# Chronic Perinatal Hypoxia Delays Cardiac Maturation in a Mouse Model for Cyanotic Congenital Heart Disease

**DOI:** 10.1101/2020.05.28.117648

**Authors:** Jennifer Romanowicz, Zaenab Dhari, Devon Guerrelli, Colm Mulvany, Marissa Reilly, Luther Swift, Nimisha Vasandani, Manelle Ramadan, Nobuyuki Ishibashi, Nikki Gillum Posnack

## Abstract

**Background:** Compared to acyanotic congenital heart disease (CHD), cyanotic CHD has an increased risk of lifelong mortality and morbidity. These adverse outcomes may be attributed to delayed cardiomyocyte maturation, since the transition from a hypoxic fetal milieu to oxygen rich postnatal environment is disrupted. We established a rodent model to replicate hypoxic myocardial conditions spanning perinatal development, and tested the hypothesis that chronic hypoxia impairs cardiac development.

**Methods:** Mouse dams were housed in hypoxia beginning at embryonic day 16. Pups stayed in hypoxia until postnatal day (P)8 when cardiac development is nearly complete. Global gene expression was quantified at P8 and at P30, after recovering in normoxia. Phenotypic testing included electrocardiogram, echocardiogram, and *ex-vivo* electrophysiology study.

**Results:** Hypoxic animals were 48% smaller than controls. Gene expression was grossly altered by hypoxia at P8 (1427 genes affected), but normalized after recovery (P30). Electrocardiograms revealed bradycardia and slowed conduction velocity in hypoxic animals at P8, which resolved after recovery (P30). Notable differences that persisted after recovery (P30) included a 65% prolongation in ventricular effective refractory period, sinus node dysfunction, and a 24% reduction in contractile function in animals exposed to hypoxia.

**Conclusions:** We investigated the impact of chronic hypoxia on the developing heart. Perinatal hypoxia was associated with changes in gene expression and cardiac function. Persistent changes to the electrophysiologic substrate and contractile function warrant further investigation, and may contribute to adverse outcomes observed in the cyanotic CHD population.

## Introduction

Outcomes for patients with cyanotic congenital heart disease (CHD) remain guarded despite countless advances in clinical care strategies over the last decades. CHD affects 1% of live births^1^, with one quarter of CHD representing cyanotic conditions^2^. Infants with cyanotic CHD are at an 8-fold increased risk of death^2^ compared to acyanotic CHD, and up to 39% of all patients with CHD develop heart failure during childhood^3^. With clinical advances, the CHD population is increasingly surviving to adulthood, leading to an increased burden of CHD-associated heart failure^4^. Despite the high incidence of morbidity and mortality in the cyanotic CHD population, the underlying risk factors are not fully understood^3^.

Normal embryology of the heart includes streaming of the most highly saturated blood to the ascending aorta which directly supplies the head vessels and coronary arteries without mixing with desaturated blood from the ductus arteriosus^5,6^. For a fetus with complex CHD—such as hypoplastic left heart syndrome or D-looped transposition of the great arteries—this streaming pattern is disrupted and the blood in the ascending aorta is desaturated compared to the normal fetal circulation^7–10^. This relative desaturation of ascending aortic blood leads to chronic hypoxic conditions in the developing brain and heart starting prenatally and persisting after birth until definitive repair. At the time of cardiac surgery, the cyanotic myocardium has depleted endogenous antioxidants^11^, higher tissue lactate levels^12^, more troponin I release^13^, higher levels of oxidative stress^14^, and less available adenosine triphosphate (ATP)^15^ compared to acyanotic myocardium. Moreover, cyanotic infants exhibit more myocardial injury during bypass surgery and worse postoperative outcomes^13,15^. Little is known about the mechanisms that contribute to the cyanotic myocardium’s vulnerability to metabolic derangements during surgery.

The transition from the hypoxic fetal milieu to the oxygen-rich postnatal environment is thought to stimulate postnatal maturation^16^. Accordingly, a limited number of studies suggest that hypoxia delays cardiac maturation. In mice, postnatal hypoxia prolongs the neonatal period of cardiomyocyte proliferative ability^17^; and in chickens, prenatal hypoxia results in immature calcium handling^18^. Moreover, cardiomyocytes sampled from human patients with hypoplastic left heart syndrome show some persistence in fetal gene programming^19^. However, the direct effects of chronic perinatal hypoxia on the developmental processes of the cardiomyocyte remain largely unknown.

To the best of our knowledge, this is the first study to examine the combined effects of pre- and postnatal hypoxia on the developing heart. The current study aimed to establish a rodent model of chronic perinatal hypoxia, as would be seen in cyanotic CHD, to investigate the developmental status of the cardiomyocyte under these conditions. We hypothesized that exposure to chronic hypoxia, beginning prenatally and continuing through the neonatal period, would perturb cardiomyocyte gene expression, contractile function, and the electrophysiologic substrate of the heart. The Cardiac Safety Research Consortium has implored the research community to perform more studies of developmental cardiac physiology to better understand the substrate on which therapies may work in the pediatric population^20^, and this study intended to contribute to that call for knowledge.

## Methods

### Disclosure Statement and Ethical Approval

Data that support the findings of this study are available from the corresponding author upon reasonable request. Animal experiments were approved by the Institutional Animal Care and Use Committee at Children’s National Research Institute, in compliance with the *NIH Guide for the Care and Use of Laboratory Animals*.

### Animal Model

Wild-type pregnant CD1 mouse dams (6-8-week-old Crl:CD1(ICR) mice, Charles River Laboratories) were kept in a hypoxic chamber (BioSpherix, Redfield, NY) starting on embryonic day (E)16, a time that coincides with myocardial reliance on coronary flow for oxygen delivery and the beginning of a period of rapid growth of the ventricular myocardium, similar to the second trimester in human fetuses^21,22^ **(Figure 1)**. Blood in the ascending aorta is desaturated in complex CHD^7^, leading to desaturated coronary arterial flow and hypoxic myocardial conditions once the myocardium is dependent on the coronary arteries for oxygenation. During the experiment, the oxygen concentration was maintained, monitored, and recorded continuously with sensors placed inside the chamber to achieve a level of 10.5 ± 0.5% (Pro:Ox Model 360, BioSpherix, Redfield, New York). Nitrogen gas was used to displace oxygen. Dams gave birth in the hypoxic chamber and pups remained in hypoxia until postnatal day (P)8, when cardiomyocyte maturation is nearly complete^23,24^. Strain and age-matched normoxic dams were kept in normoxia and gave birth under normoxic conditions. After P8, hypoxic animals were moved to normoxic conditions and allowed to recover until further testing at P30, thus simulating elevated oxygen saturations in human infants who have undergone definitive repair of cyanotic CHD. At the end of study, animals were euthanized via carbon dioxide inhalation and cervical dislocation. For ex vivo electrophysiology studies, animals were anesthetized with 4% isoflurane, the heart was excised, and euthanasia ensued via exsanguination.

**Figure 1:**
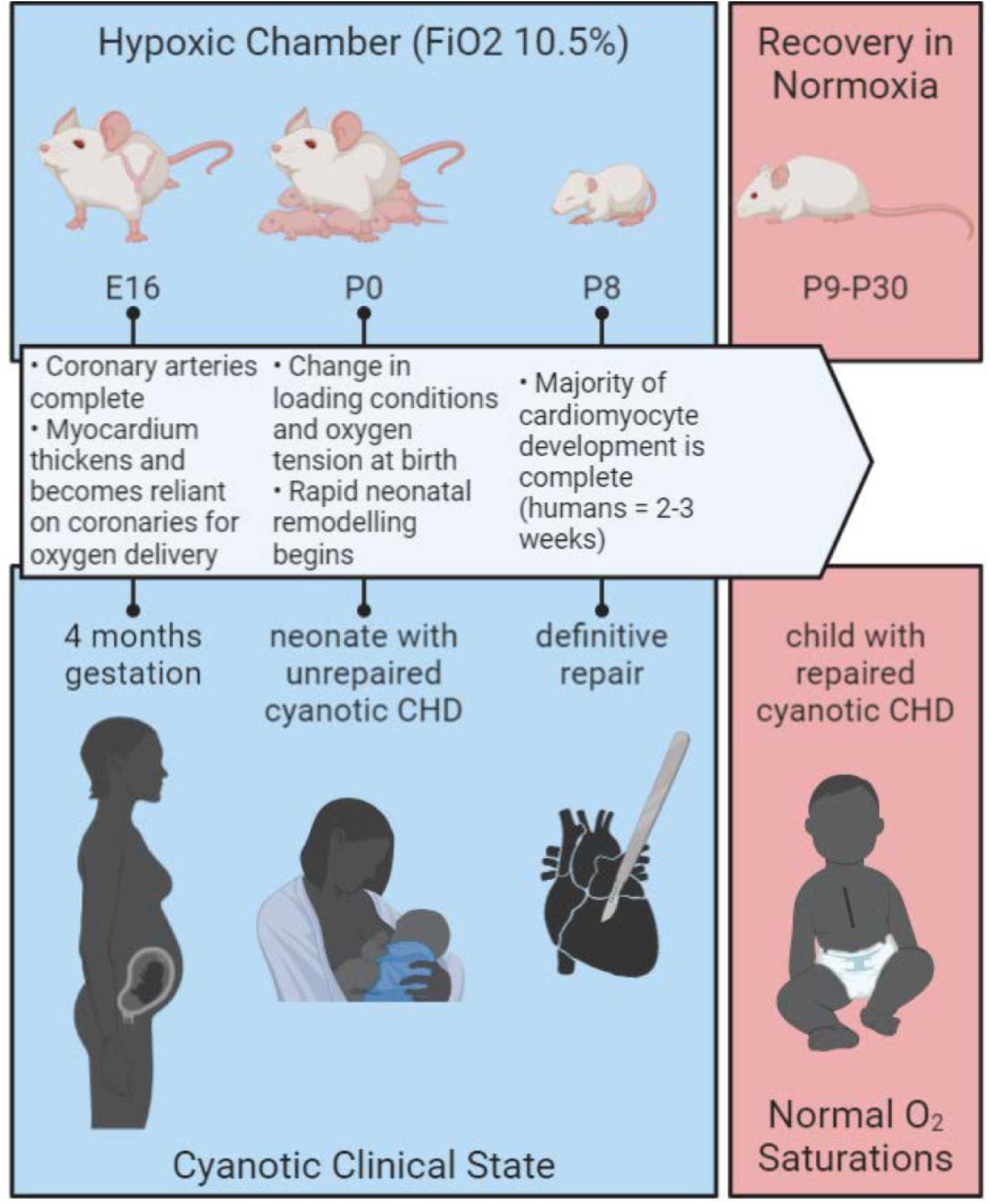
Chronic Perinatal Hypoxia Mouse Model. Pregnant CD1 mice were placed in hypoxia starting on embryonic day (E)16, corresponding with rapid growth of the ventricular myocardium and reliance on coronary arteries, similar to a 4 months gestation human fetus. Hypoxic pups were born and reared in hypoxia until postnatal day (P)8 when the majority of cardiomyocyte development is complete. Removal from the hypoxic chamber represents definitive repair in human neonates with cyanotic CHD. Mice recovered in normoxia until further testing at P30, representing a child with repaired cyanotic CHD and corresponding normalization in oxygen saturations. FiO2, fractional inspired oxygen concentration. Image created with Biorender.com.

### Gene Expression

Whole hearts were excised and flash frozen in liquid nitrogen. Total RNA was isolated using an RNeasy fibrous tissue kit. Verification of RNA integrity and RNA quantification were done by spectrophotometry and an RNA 6000 Nano assay (Bioanalyzer 2100, Agilent Technologies). The RNA integrity number for all samples was >6 (mean=8.6 +/-SEM 0.19). Total RNA (250 ng) was primed for the entire length of RNA, including both poly(A) and non-poly(A) mRNA and reverse transcribed to generate sense-strand targets that were biotin-labelled using a WT Plus Reagent kit, and then hybridized to Affymetrix GeneChip Mouse Clariom S arrays for 16 hours (48°C), following manufacturer’s instructions (Thermo Fisher Scientific). Hybridization cocktails were removed, and arrays were washed and stained on a Fluidics Station 450 (mouse Clariom S arrays). Arrays were scanned on the Affymetrix GCS3000 7G scanner and initial quality control data evaluated using Affymetrix Expression Console software (Thermo Fisher Scientific). Microarray data was imported and analyzed (ANOVA p<0.05, 1.5 fold cutoff, 0.1 false discovery rate) using the Transcriptome Analysis Console (Applied Biosciences). Gene ontology (GO) enrichment analysis was performed with the GOrilla tool using a single rank-ordered gene list^25,26^.

### In-Vivo Electrocardiography

Non-invasive electrocardiogram (ECG) recordings were obtained using an ecgTUNNEL system (emka Technologies). ECG waveforms were recorded for two minutes on P8 conscious and isoflurane-sedated animals as well as P30 isoflurane-sedated animals (litter mates). ECG segments were quantified using ecgAuto software (emka Technologies) for heart rate, heart rate variability, atrial depolarization time (P-wave duration), atrioventricular conduction time (PR interval), ventricular depolarization time (QRS duration), and ventricular repolarization time (QT interval; uncorrected as heart rate does not significantly alter QT interval in mice^27^). Heart rate variability was measured as a root mean square of the successive differences (RMSSD)^28,29^. Due to motion artifact, only clearly discernable ECG parameters were included in the analysis.

### Ex-Vivo Electrophysiology Study

P30 animals were anesthetized with 4% isoflurane, the heart was rapidly excised and the aorta cannulated. The heart was transferred to a temperature-controlled (37°C) constant-pressure (70 mmHg) Langendorff perfusion system. Excised hearts were perfused with Krebs-Henseleit buffer bubbled with carbogen, as previously described^29,30^. A stimulation electrode was placed externally on the right atrium, and an atrial pacing protocol was used to determine Wenckebach cycle length (WBCL) and atrioventricular nodal effective refractory period (AVNERP). WBCL is the shortest pacing cycle length during atrial pacing that causes the Wenckebach phenomenon. AVNERP is the shortest extrastimulus interval during atrial pacing that fails to conduct through the atrioventricular (AV) node, as indicated by loss of ventricular capture. For ventricular pacing, a stimulation electrode was placed on the LV epicardium. To determine the ventricular effective refractory period (VERP), dynamic pacing was performed with stepwise decrements in the pacing cycle length (S1-S2) until loss of capture was noted. Baseline rhythms were monitored throughout the duration of the studies for detection of dysrhythmias including ectopy, sinus node dysfunction, and AV nodal block. Sinus node dysfunction was defined as bradycardia with an irregular sinus rate.

### High Frequency Ultrasound Echocardiography

P30 animals underwent sedated transthoracic echocardiography to assess the persistent effects of chronic perinatal hypoxia on left ventricular systolic function. Anesthesia was initiated and maintained with inhaled isoflurane (1.5-2%). A pre-clinical high-frequency ultrasound system (VisualSonics Vevo 770, 30mHz probe) was used to obtain fractional shortening in a parasternal short axis view using M-mode measurements. Measurements of interventricular septum and left ventricular posterior wall thickness were measured on the same M-mode images.

### Statistical Analysis

Statistical analysis was performed using GraphPad Prism. Data normality was confirmed by Shapiro-Wilk test. Datasets were compared between control and hypoxic animals (biological replicates) using two-tailed t-test or ANOVA with 0.1 false discovery rate to control for multiple comparisons testing (q value reported). Significance was defined as *p<0.05.

## Results

### Hypoxia decreased litter size and pup weight

Hypoxic litters had markedly fewer pups than normoxic litters (n≥5 litters per group, mean 5.7 vs 12.4 pups per litter, p=0.0005, **Figure 2A**). Hypoxic animals were smaller than normoxic controls at P8 (n≥25, mean 3.4 vs 6.5 g, p<0.0001, **Figure 2B-C**). Despite lower body weight, heart weight was preserved in hypoxia such that it did not differ from control (n=10, mean 46 vs 44 mg, p=0.58). As a result, heart-to-body-weight ratios were higher in hypoxic animals (n=10, mean 8.0 vs 15.4 mg/g, p<0.0001). After P8, hypoxic animals recovered in normoxic conditions, thus simulating elevated oxygen saturations that occurs after definitive repair of cyanotic CHD. At P30, hypoxic animals had undergone catch-up growth such that their body weight was only slightly lower than normoxic controls (n≥14, mean 20.0 vs 22.7 g, p=0.015, **Figure 2D**). Heart weight trended toward slightly lower in hypoxic animals compared to control (n≥9, mean 119 vs 141 mg, p=0.079), such that heart-to-body-weight ratios were equivalent between the groups (n≥9, mean 6.4 vs 6.3 mg/g, p=0.95).

**Figure 2:**
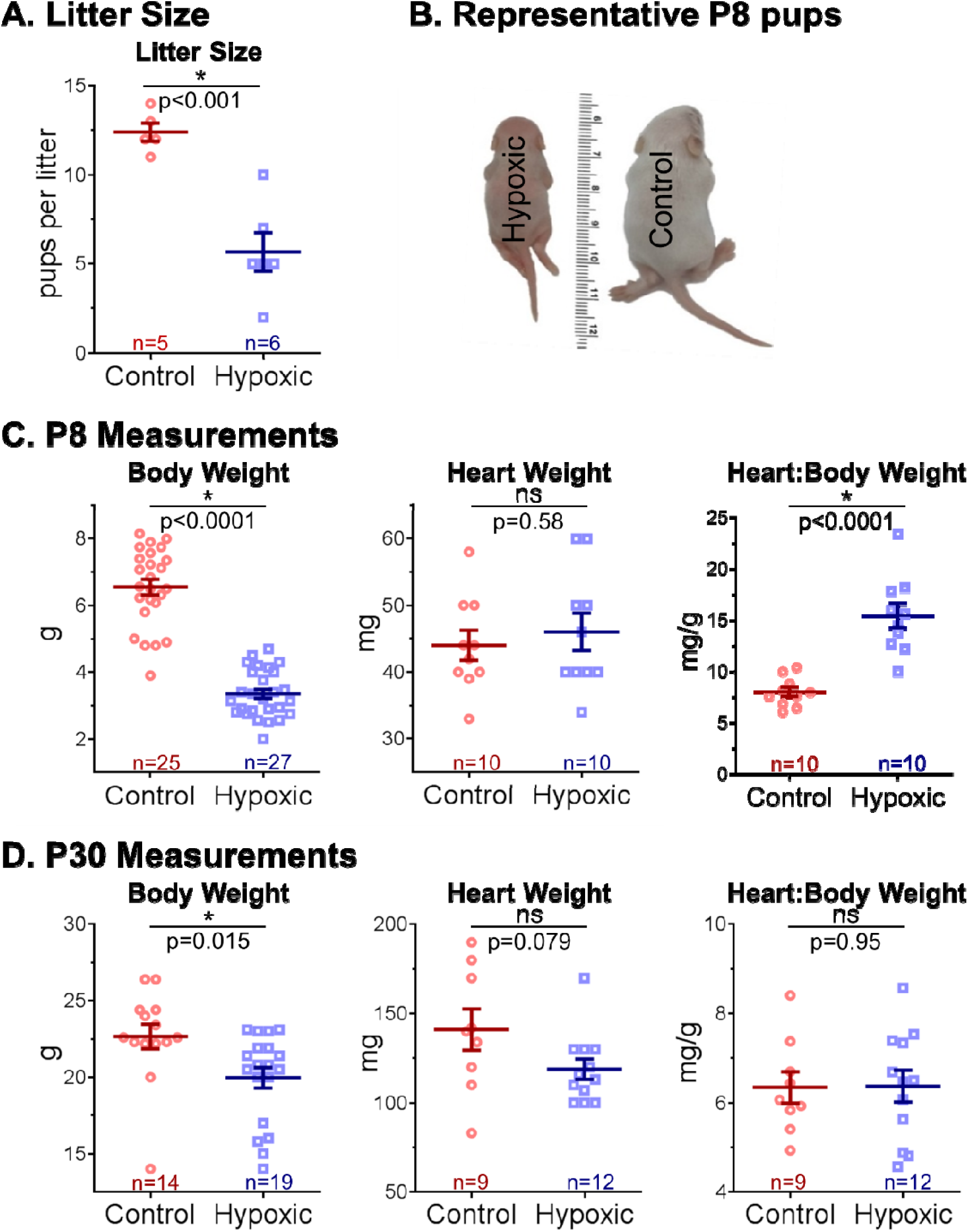
Litter Size and Body Measurements. **A:**Hypoxia reduced mean litter size. **B, C**: At P8, hypoxic animals were markedly smaller than control; however, heart weight was preserved, and thus heart-to-body-weight ratio was higher in hypoxic animals. **D:**By P30, hypoxic animals underwent catch-up growth such that they were only slightly smaller than control; heart weight trended toward slightly lower in hypoxic animals, and heart-to-body weight ratios were equivalent between groups. Biological replicates are shown and sample size indicated. Data expressed as mean ± SEM. *p<0.05 hypoxic versus control via two-tailed t-test, ns=not significant.

### Hypoxia altered global gene expression at P8

Gene expression arrays were performed on whole heart samples isolated at P8 to assess the effects of hypoxia at the end of the neonatal period of rapid cardiomyocyte development, and again at P30 after hypoxic animals had recovered in normoxia. Principal component analysis demonstrated that experimental groups (normoxia versus hypoxia) were well-separated by their mRNA expression profiles at P8, but this separation was negligible at P30 (**Figure 3A**). Using a 1.5-fold expression cut-off and a 10% false discovery rate to correct for multiple testing^31^, a total of 1427 mRNAs were differentially expressed between hypoxic and control hearts at P8 (**Figure 3B**; data available via Gene Expression Omnibus). Differentially-expressed genes important to cardiac functioning and development are highlighted in the **Figure 3C**volcano plot. Within treatment groups, hypoxic animals exhibited a greater number of gene changes between P8 and P30 (4593 genes; **Figure 3B**) than the normoxic group (2147 genes), suggesting that the hypoxic group underwent more developmental changes over this time period after transitioning to normoxia. Interestingly, gene expression nearly normalized in P30 animals after recovering in normoxia. Only one gene was expressed differentially between groups at P30: Wsb2 (WD repeat and SOCS box-containing 2).

**Figure 3:**
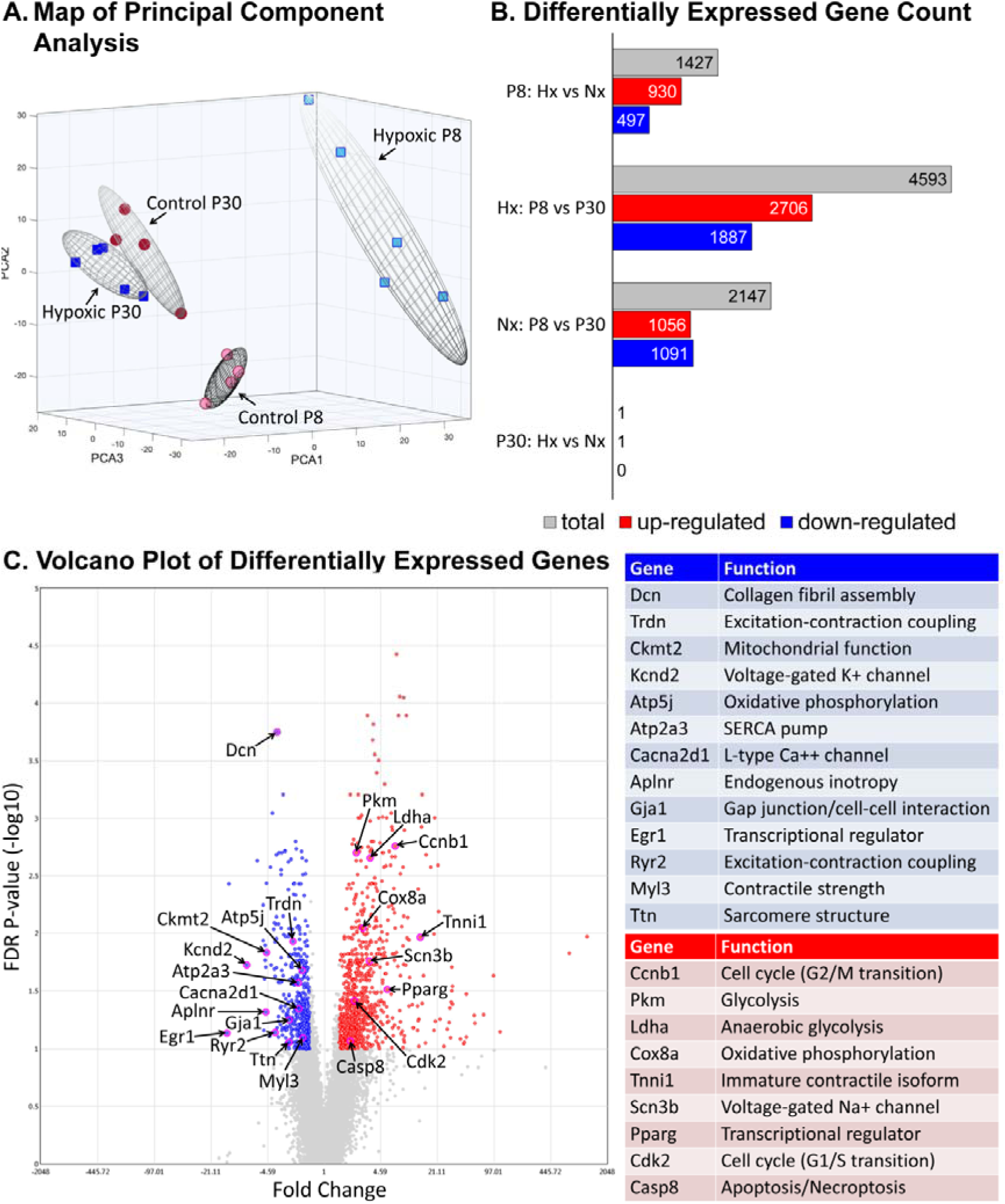
Global Gene Expression. **A:**Principal component analysis demonstrated that experimental groups were well separated by their mRNA expression profiles at P8 (n=5 hypoxic, n=4 control biological replicates), but this separation was negligible in P30 samples (n=5 hypoxic, control biological replicates). **B:**A total of 1427 mRNAs were differentially expressed between hypoxic (Hx) and normoxic control (Nx) hearts at P8, with near resolution of differences by P30 (ANOVA with 1.5-fold expression cut-off, p<0.05 and false-discovery rate q≤0.1). **C:**Specific genes important to cardiovascular functioning and development are highlighted in a volcano plot of all differentially expressed genes between groups at P8.

Differentially expressed genes at P8 were significantly overrepresented in >400 GO categories (319 biological processes, 42 molecular functions, 46 cellular components; data available via Gene Expression Omnibus). Categories associated with phenotypic changes observed in our experimental studies included extracellular matrix structural constituent (GO:0005201), ion channel binding (GO:0044325), glycolytic process (GO:0006096), hypoxia-inducible factor-1alpha signaling pathway (GO:0097411), mitotic cell cycle process (GO:1903047), and cell maturation (GO:0048469) (**Figure 4**).

**Figure 4:**
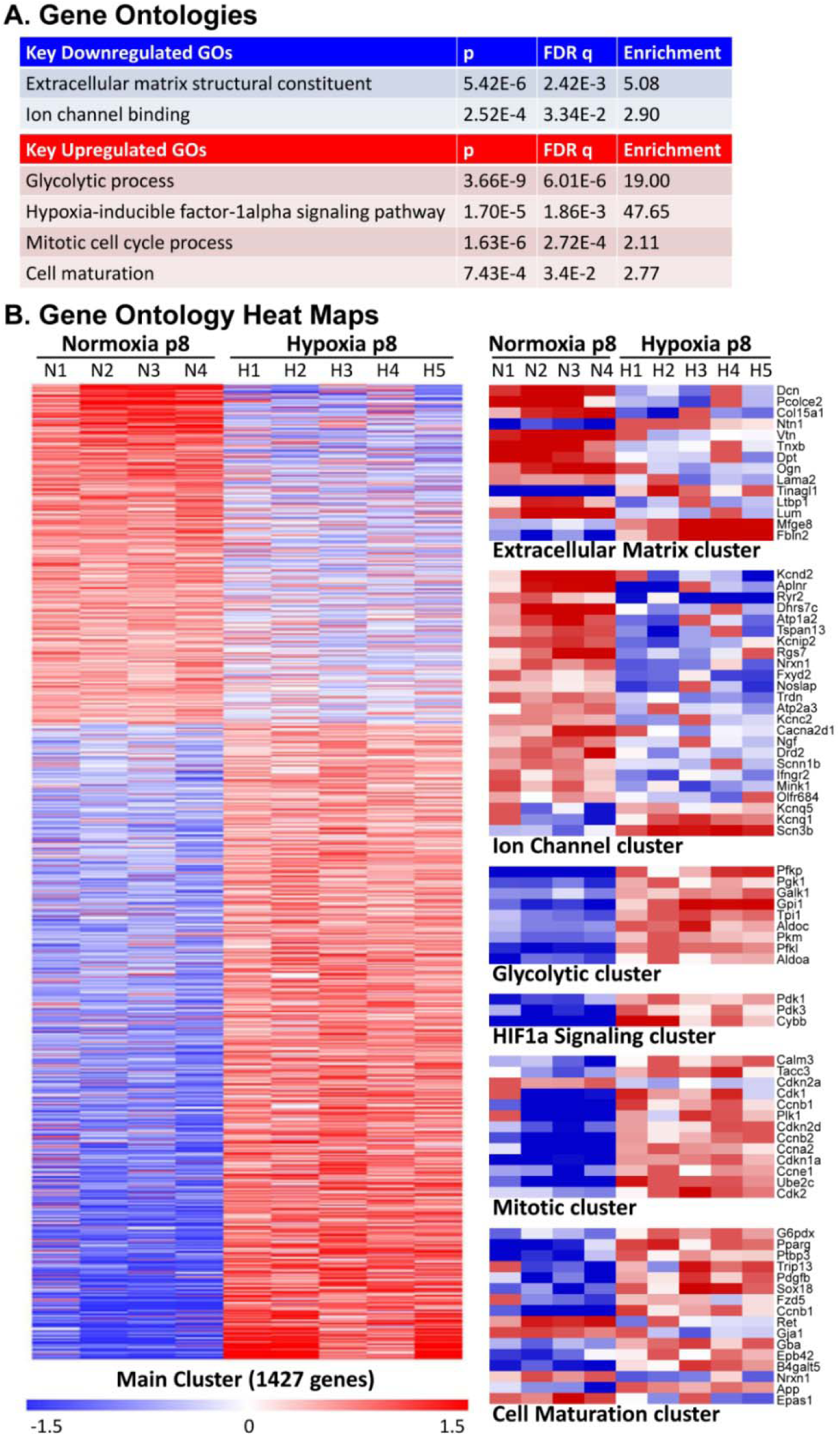
Gene Ontologies. **A:**Highlighted gene ontologies (GOs) important to cardiac development and functioning that were different between groups at P8 (see Figure 3). GOs based on differential gene expression between hypoxic (Hx) and normoxic control (Nx) hearts at p8 (ANOVA, p<0.05, false-discovery rate q≤0.1).**B:**Heat maps of the main cluster of differentially expressed genes, as well as selected gene ontologies demonstrate differential expression at P8 between groups. Each gene is median-centered, with data displayed as fold-change. n=5 hypoxic, n=4 control biological replicates.

### Hypoxia altered transmembrane ion channel expression at P8

Multiple transmembrane ion channels were differentially expressed in P8 hypoxic animals, including potassium (K^+^), sodium (Na^+^), and calcium (Ca^++^) channels involved in the cardiac action potential and maintenance of a stable resting membrane (**Figure 5A**). In ventricular myocytes, phase 0 of the cardiac action potential is characterized by rapid depolarization via Na^+^ influx (I_Na_); voltage-gated Na^+^ channel gene expression was upregulated with hypoxia (Scn3b: fold change +3.4, p=0.0002, q=0.017; Scn1b: fold change +1.3, p=0.018, q=0.15). Phase 1 is characterized by a fast transient outward K^+^ current (I_to_), which was downregulated in hypoxia (Kcnd2: fold change −8.02, p=0.0003, q=0.019; Kcnip2: fold change −3.17, p=0.0012, q=0.042; Kcnd3: fold change −1.46, p=0.0079, q=0.10). Notably, decreased I_to_ current is also observed in patients with atrial fibrillation^32^ and heart failure^33^, and can impair electromechanical coupling and prolong action potential duration^34^. Phase 1 also includes slow Na^++^ efflux via the Na^+^/Ca^++^ exchanger (NCX1), which was upregulated in hypoxia (Slc8a1: fold change +2.78, p=0.0001, q=0.012). The plateau phase (phase 2) is primarily responsible for action potential duration. Membrane potential is held stable by balancing Ca^++^ influx and K^+^ efflux, and both were affected by hypoxia. Ca^++^ influx continues via NCX1 (upregulated in hypoxia, see above), and L-type voltage-gated Ca^++^ channels open to increase Ca^++^ influx (I_CaL_). I_CaL_ channels were downregulated in hypoxia (Cacna2d1: fold change −1.96, p=0.0014, q=0.045). Calmodulin expression was increased (Calm3: fold change +1.67, p=0.0009, q=0.036) which modulates both action potential duration and excitation-contraction coupling by modifying I_CaL_. Slow K^+^ efflux (I_Ks_) occurs via voltage-gated K^+^ channels, which were upregulated in hypoxia (Kcnq1: fold change +2.54, p=0.0057, q=0.089). Final rapid repolarization (phase 3) occurs mainly by K^+^ efflux (I_Kr_). Both genes associated with I_Kr_ trended toward downregulation (Kcne2: fold change −1.47, p=0.22, q=0.51; Kcnh2: fold change −1.24, p=0.28, q=0.57). Finally, phase 4 represents the resting membrane potential which is maintained via constant K^+^ efflux (I_K1_, I_Ach_, I_ATP_) through inwardly rectifying K^+^ channels; these genes were largely unaffected by hypoxia.

**Figure 5:**
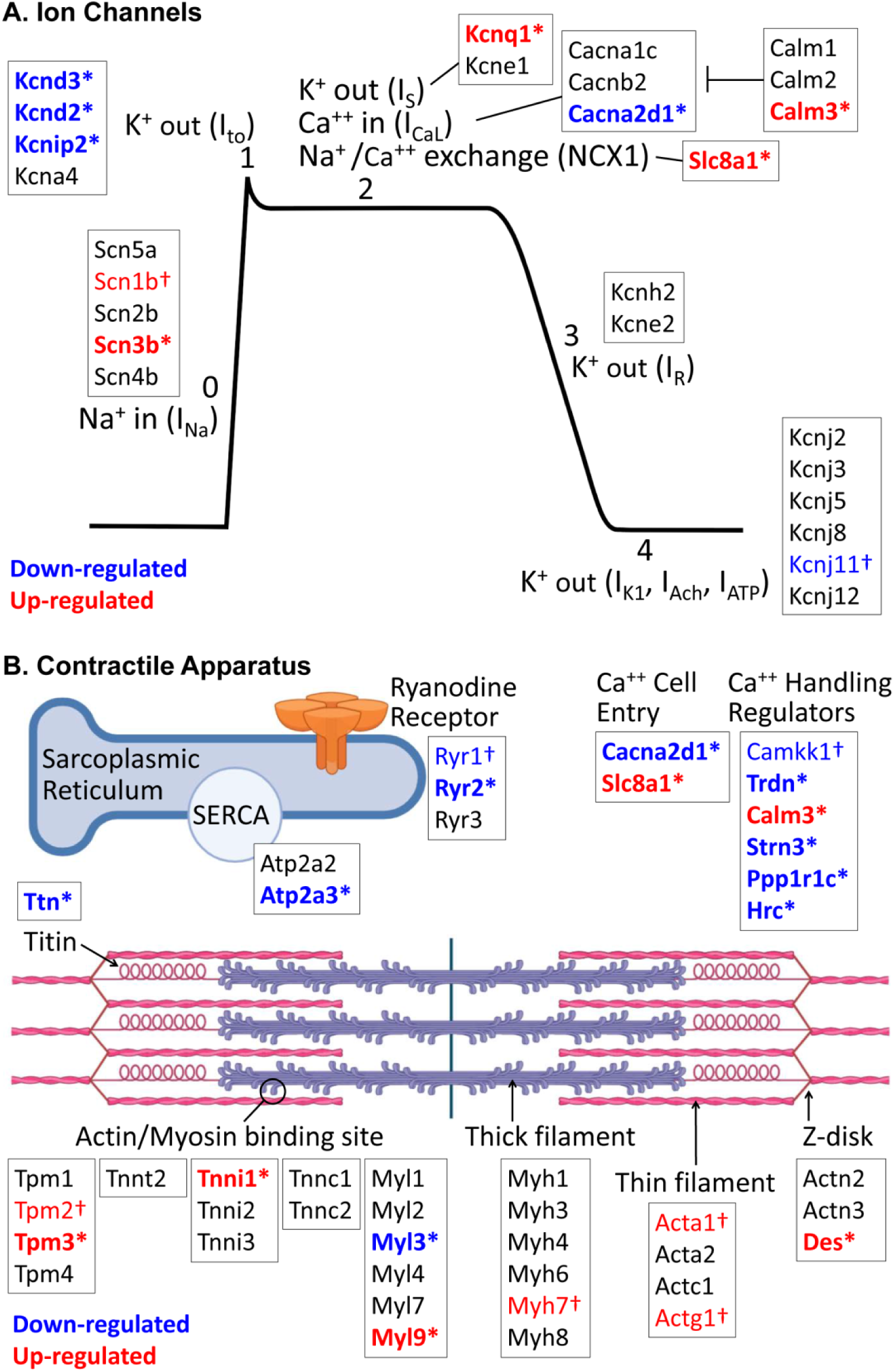
Genes Affecting Ion Channels and the Contractile Apparatus. **A:**At P8, hypoxia affected genes involved with most phases of the cardiac action potential. **B:**Hypoxia affected multiple genes important to the contractile apparatus, including both the sarcomere and calcium handling machinery, at P8. Results based on differential gene expression. Significance denoted by *p≤0.05 and FDR q≤0.1, or †p≤0.05 and FDR q>0.1. n=5 hypoxic, n=4 control biological replicates. Image created with Biorender.com.

In cardiac pacemaker cells, the T-type Ca^++^ channel generates Ca^++^ influx (I_CaT_) to initiate an action potential (gene = Cacna1g). Cacna1g had a 3.13 fold down-regulation which did not meet significance (p=0.052, q=0.26). Additional studies are needed to examine regional differences in T-type Ca^++^ channel expression, which can result in bradycardia and sinus node dysfunction if localized to pacemaker cells. Finally, gap junctions connect neighboring cardiomyocytes and allow for rapid spread of an action potential from one cell to the next. Gap junction expression was downregulated in hypoxia (Gja1: fold change −2.57, p=0.0022, q=0.056; Gja6: fold change − 1.74, p=0.0067, q=0.096), which can lead to slower electrical conduction.

### Hypoxia altered expression of genes important to the contractile apparatus at P8

Multiple genes important to the contractile apparatus were differentially expressed in P8 hypoxic animals, including those involved in both calcium handling and sarcomere structure (**Figure 5B**). Within the cardiac sarcomere, isoform switching occurs during perinatal development for many key structural proteins. In hypoxic P8 animals, there was persistence of immature isoforms of Troponin-I (Tnni1: fold change +13.13, p=9.71E-05, q=0.011), alpha-actin (Acta1: fold change +5.57, p=0.012, q=0.12), gamma-actin (Actg1: fold-change +3.05, p=0.010, q=0.12), and myosin heavy chain (Myh7: fold change +1.48, p=0.083, q=0.32); although only Troponin-I met the predetermined threshold for significance. The isoform switch for myosin light chain was unaffected (Myl7→Myl2), however expression of two myosin light chain regulators were altered (Myl3: fold change −1.79, p=0.0045, q=0.08; Myl9: fold change +5.48, p=0.0025, q=0.06).

Multiple stabilizing components of the cardiac sarcomere were also altered in P8 hypoxic animals. Titin, the cardiac myofilament responsible for passive tension, was downregulated in hypoxia (fold change −2.56, p=0.0055, q=0.088), which may cause an increased risk for diastolic dysfunction^35^. Desmin provides strength of attachment of myofibrils to the Z-disk, and was upregulated in hypoxia (Des: fold change +2.31, p=0.0023, q=0.057). Desmosomes and adherens junctions provide structural integrity between cells, and both demonstrated downregulation of components in hypoxia (Pkp1: fold change −1.55, p=0.0022, q=0.056; Pcdh7: fold change −1.82, p=0.0025, q=0.061; Dsc2: fold change −2.04, p=0.016, q=0.14). Vimentin expression was increased (Vim: fold change +3.08, p=0.0017, q=0.050) indicating an increased fibroblast population, but collagen expression was decreased (extracellular matrix cluster in **Figure 4B**), suggesting decreased fibroblast functioning and a less robust extracellular matrix to act as a scaffold for muscle contraction.

Calcium handling genes were also altered in hypoxic P8 animals. In mature cardiomyocytes, synchronized ryanodine receptors facilitate a rapid increase in cytosolic Ca^++^ concentration which is necessary for excitation-contraction coupling. Ryanodine receptor expression was decreased in hypoxic animals (Ryr2: fold change −3.74, p=0.0038, q=0.073; Ryr1: fold change −1.16, p=0.029, q=0.20), consistent with a delay in maturation. Further, extracellular Ca^++^ entry may be altered in hypoxic animals due to decreased expression of L-type calcium channels (Cacna2d1, see above) and increased expression of the Na^+^/Ca^++^ exchanger (Slc8a1, see above). The sarco-endoplasmic reticulum Ca^++^-ATPase (SERCA) pumps calcium from the cytoplasm back into the sarcoplasmic reticulum in preparation for the next cardiac cycle. Hypoxia decreased SERCA expression (Atp2a3: fold change −2.03, p=0.0005, q=0.027), which could decrease sarcoplasmic reticulum Ca^++^ load, reduce contractility, and slow lusitropy.

### Hypoxia caused bradycardia and slowed conduction at P8

With the observed changes in ion channel expression on gene arrays, we collected *in-vivo* ECG tracings at P8 (**Figure 6A**) to identify alterations in the electrophysiologic substrate of the heart immediately after chronic perinatal hypoxia. During normal murine development, heart rate increases as maturation progresses. At P8, hypoxic animals were bradycardic compared to normoxic controls (n≥10, mean 306 vs 562 beats per minute, q<0.0001), consistent with a delay in normal postnatal maturation (**Figure 6B**). We examined the effects of isoflurane sedation on ECG measurements, as cyanotic newborns with CHD are subjected to anesthesia at the time of surgical repair. Hypoxic animals remained more bradycardic than normoxic when sedated (n≥10, mean 204 vs 272 beats per minute, q=0.003, **Figure 6B**).

**Figure 6:**
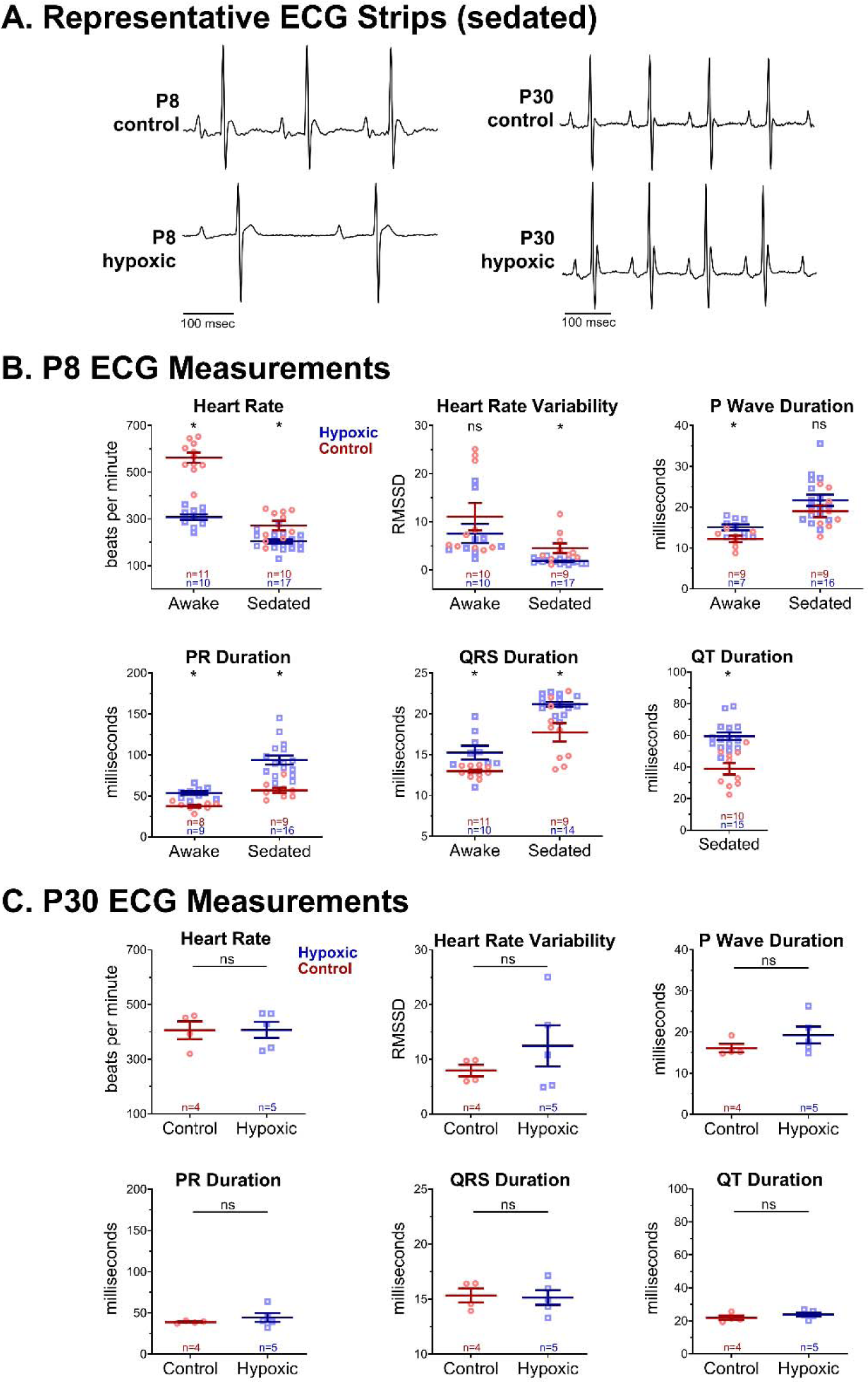
In-Vivo Electrocardiogram Measurements. **A:**Representative ECG traces (isoflurane-sedation shown) **B:**Awake hypoxic P8 animals had lower heart rates compared to control, and hypoxia prolonged ECG intervals, consistent with slowed conduction speed. Sedated hypoxic P8 animals exhibited slower heart rate, lower heart rate variability, and conduction slowing in most ECG parameters compared with control. **C:**At P30, all ECG measurements had normalized under isoflurane-sedation. Biological replicates are shown and sample size indicated. Data expressed as mean ± SEM. *p<0.05 hypoxic versus control via two-tailed t-test, ns=not significant. Due to motion artifact, only clearly discernable ECG parameters were included in the analysis.

Conduction speed is expected to increase throughout development as cell-cell interactions mature. Awake measurements indicated that P8 hypoxic animals had a longer P-wave duration (n≥7, mean 15.0 vs 12.2 msec, q=0.025), PR interval (n≥8, mean 53.4 vs 37.5 msec, q=0.0001), and QRS duration (n≥10, mean 15.3 vs 13.0 msec, q=0.01). In vivo ECG parameters were also altered under isoflurane sedation: compared with control, hypoxic P8 animals exhibited reduced heart rate variability (n≥10, mean 1.8 vs 4.5, q=0.002) as well as longer QRS durations (n≥10, mean 21.1 vs 17.7 msec, q=0.005), PR intervals (n≥10, mean 93.8 vs 56.7 msec, q=0.0001), and QT intervals (n≥10, mean 59.4 vs 38.8 msec, q<0.0001) (**Figure 6B**). For accuracy, QT intervals were only analyzed in sedated animals. Anesthetic agents have been reported to slow atrioventricular conduction in animal models^36,37^ and human case reports^38,39^, and our results suggest that perinatal hypoxia may exaggerate this effect. We also observed a significantly lower heart rate variability in hypoxic animals compared with normoxic controls, and heart rate variability increases with age during normal development^29^.

### ECG measurements normalized at P30 after a period of recovery in normoxia

Since there was resolution of genetic differences at P30 after recovering in normoxia, ECG measurements were repeated at P30 to determine if electrophysiologic differences had also normalized. Under isoflurane sedation, we observed no significant difference in heart rate (n≥4, mean 407 vs 406 beats per minute, p=0.98), heart rate variability (n=4, mean 12.5 vs 8.0 msec, p=0.34), P-wave duration (n≥4, mean 17.7 vs 15.2 msec, p=0.28), PR interval (n≥4, mean 44.4 vs 38.9 milliseconds, p=0.39), QRS duration (n≥4, mean 15.1 vs 15.3 msec, p=0.84) and QT interval (n≥4, mean 23.9 vs 21.9 msec, p=0.29) between groups at P30 (**Figure 6C**). The significant differences in ECG parameters observed at P8 largely abated after the 22 day period of recovery in normoxia, consistent with gene expression data (**Figure 3A-B**).

### Ex-vivo electrophysiology study revealed persistent underlying changes to the electrophysiologic substrate after hypoxia

Although ECG measurements largely normalized by P30, we conducted more rigorous testing of the cardiac electrophysiologic substrate in the absence of autonomic influences. Electrophysiology studies performed at P30 revealed that perinatal hypoxia caused persistent prolongation of VERP compared to normoxic controls (n=4, mean 76.5 vs 46.5 msec, p=0.013), a parameter that normally decreases with age (**Figure 7A**). No difference in atrioventricular conduction was observed between groups, measured by WBCL (n=4, mean 84.0 vs 84.5 msec, p=0.94) and AVNERP (n=4, mean 72.8 vs 70.5 msec, p=0.76) (**Figure 7A**). *Ex-vivo* studies also revealed sinus node dysfunction in all four of the hypoxic hearts, as opposed to 25% (1/4) of the normoxic control hearts: *X*^2^(1, n=4) = 4.80, p =0.029 (**Figure 7B,C**). Cyanotic CHD carries a high incidence of sinus node dysfunction^40,41^, and our results suggest that hypoxia may play a role in creating the substrate for sinus node dysfunction.

**Figure 7:**
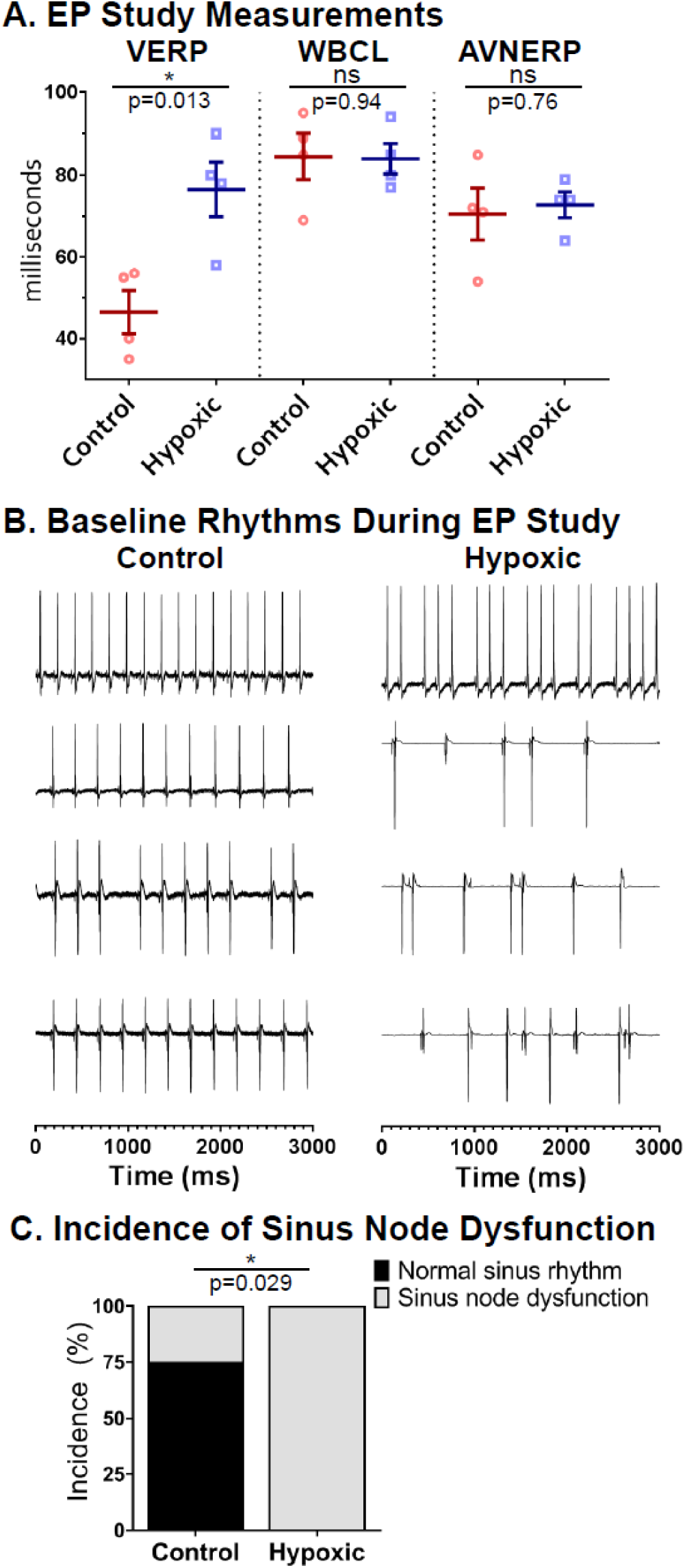
Ex-Vivo Electrophysiology Study. **A:**VERP was prolonged at P30 in animals exposed to hypoxia. There was no difference for WBCL or AVNERP. Data expressed as mean ± SEM. *p<0.05 hypoxic (n=4) versus control (n=4) via two-tailed t-test, ns=not significant. **B:**Baseline rhythm during electrophysiology studies is displayed for each animal. **B, C:**100% of hypoxic and 25% of control animals had sinus node dysfunction during the study; significance assessed using chi-squared analysis (n=4 hypoxic, control). VERP, ventricular effective refractory period. WBCL, Wenckebach cycle length. AVNERP, atrioventricular nodal effective refractory period. *p<0.05. ns=not significant. Biological replicates are shown.

### Perinatal hypoxia caused a persistent decrease in contractile function

Gene expression changes indicated differences in the sarcomere and calcium handling at P8. Although animals were too small to obtain measurements at P8, we measured phenotypic contractile function at P30 after recovery in normoxia to assess for persistent changes. Transthoracic echocardiography at P30 demonstrated decreased fractional shortening in animals exposed to hypoxia as compared to normoxic controls (n≥3, mean 0.29 vs 0.38, p=0.027, **Figure 8**), consistent with worse contractile function. There was no difference in interventricular septum thickness or left ventricular posterior wall thickness between groups at both systole and diastole.

**Figure 8:**
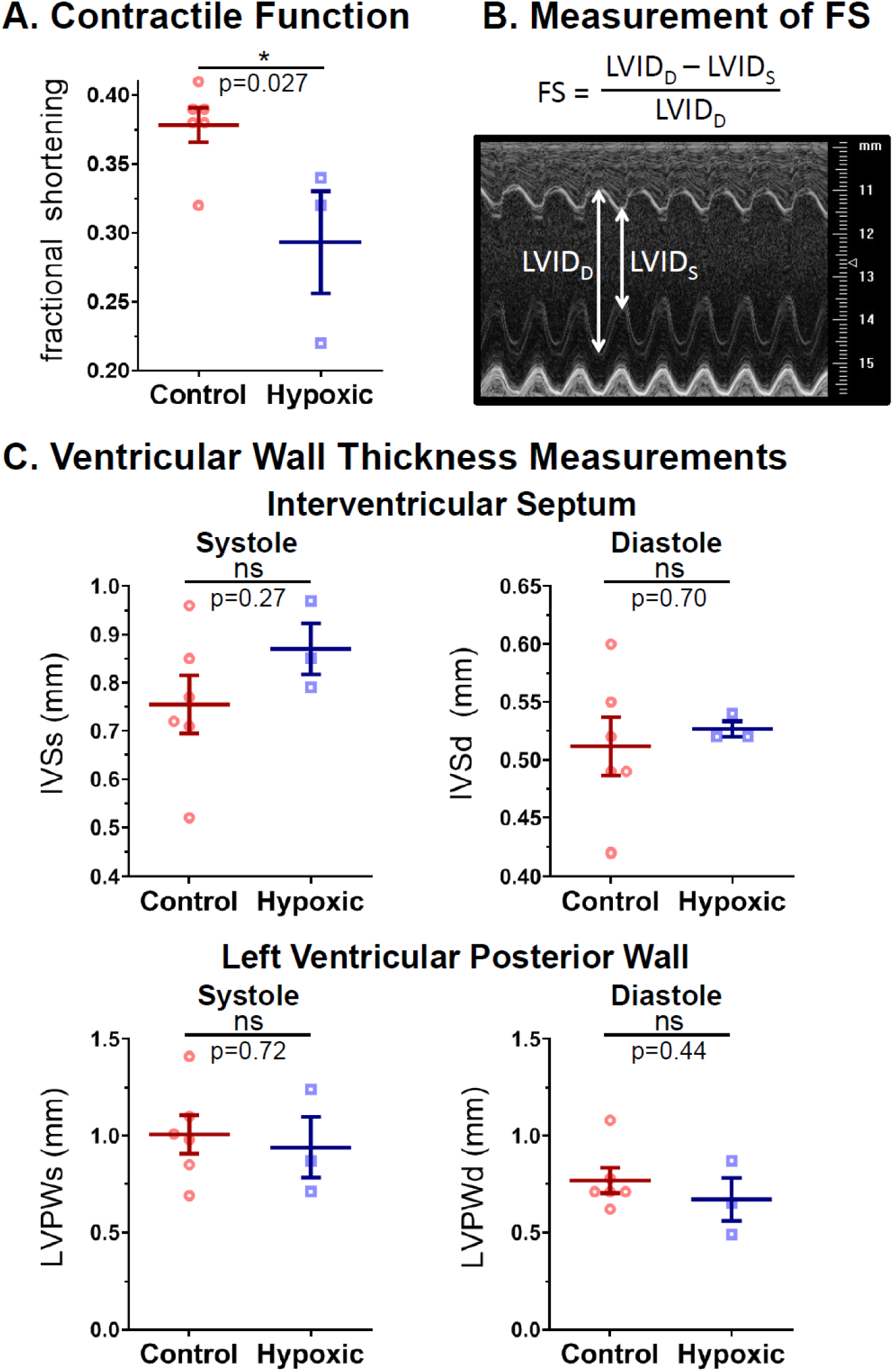
Contractile Function. **A:**Hypoxic animals had decreased contractile function compared to control, as measured by fractional shortening (FS). **B:**Measurements of the left ventricular internal diameter in systole (LVID_S_) and diastole (LVID_D_) were taken in M-Mode at the level of the papillary muscles from a parasternal short axis view. VevoSonics 770 with 30MHz high frequency ultrasound probe. **C:**There was no difference between groups in either interventricular septal thickness or left ventricular posterior wall thickness in both systole (IVSs, LVPWs) and diastole (IVSd, LVPWd). Data expressed as mean ± SEM. *p<0.05 hypoxic (n=3) versus control (n=6) via two-tailed t-test, ns=not significant. Biological replicates are shown.

## Discussion

Our results support our hypothesis that cardiac maturation is perturbed in a mouse model of chronic perinatal hypoxia. Chronic perinatal hypoxia altered both the electrophysiologic substrate and the contractile apparatus. Although the majority of differences detected at P8 normalized after recovering in normoxia, there were persistent alterations at P30 that may contribute to lifelong mortality and morbidity in the cyanotic CHD population.

Numerous genetic and phenotypic differences were detected at P8. Hypoxia altered global gene expression and ECG parameters, including bradycardia, slowed conduction speed, and decreased heart rate variability and exaggerated PR prolongation with sedation. In our animal model, P8 represents the time of surgical repair, and therefore, phenotypic differences in hypoxic animals may have implications for surgical outcomes and the immediate post-surgical course. Disturbances in ion channel expression may explain ECG disturbances at P8, and may predispose the hypoxic heart to arrhythmias. Specifically, reduced I_to_ current and an altered plateau phase can prolong the action potential, and reduced gap junctions can slow electrical conduction across the heart. Likewise, decreases in extracellular matrix collagen and alterations in the contractile apparatus have the potential to affect the strength of cardiac contraction. Electromechanical coupling development was especially delayed in hypoxic animals, as demonstrated by decreased L-type Ca^++^ channel, ryanodine receptor, and SERCA pump expression; all of which can contribute to a blunted increase in cytosolic Ca^++^ concentration and therefore weaker contraction. Increased dependence on glycolysis (**Figure 4**) may reduce myocardial energy reserves, as observed in infants undergoing surgical repair^11–15^. Furthermore, if myocardial growth continues by cell proliferation instead of hypertrophy, there is an increased risk of cell structure abnormalities. Notably, some differentially expressed genes in our study are associated with clinical sudden arrhythmic death syndromes^42^ (long QT syndrome, Brugada syndrome, arrhythmogenic right ventricular dysplasia, catecholaminergic polymorphic ventricular tachycardia) and clinical cardiomyopathies^43^ (dilated, hypertrophic, left ventricular noncompaction).

In our model, P30 represents recovery after early definitive repair of cyanotic CHD. Gene expression and ECG differences observed at P8 largely resolved by P30, suggesting that the heart was able to complete development after recovery. This is an optimistic sign that many of the observed effects from chronic perinatal hypoxia may be reversible with early repair. However, *ex-vivo* electrophysiology studies revealed persistent changes to the underlying electrophysiologic substrate including prolonged VERP and increased sinus node dysfunction, and echocardiogram revealed persistent decreased contractile function. This is similar to previous animal studies of pre- or postnatal hypoxia, which demonstrated both systolic and diastolic dysfunction^18,44–46^; however, this is the first study demonstrating persistent electrophysiologic changes after perinatal hypoxia. Importantly, measurements of gene expression do not necessarily reflect differences in protein expression and localization, ion channel current, or the myofilament and sarcomere architecture. Studies suggest that prenatal hypoxia imprints on a fetus and causes lifelong changes to the cardiovascular system, such as increased susceptibility to systemic hypertension and metabolic syndrome and worse response to myocardial infarction^47^. Further investigation is required to define proteomic, metabolic, and ion current changes that persist after recovery from chronic perinatal hypoxia.

To date, there has been limited investigation into the effects of hypoxia on the developing heart, and thus there is no established animal model. The main embryological progression of heart development is the same between humans and rodents^48^. The early embryonic heart is thin-walled and relies on diffusion of oxygen from the chambers until the coronaries connect to the aorta^48^, at which point the myocardium starts to grow and thicken^23,48^. This timeline was the rationale for starting hypoxia on embryonic day 16, when the coronary development is complete^22,49^. This timing captures the period of rapid ventricular growth that occurs in both species once the myocardium is reliant on the coronary circulation for oxygen delivery. For both humans and rodents, there are similar changes in myocyte proteins for the remainder of gestation, and both demonstrate rapid and marked development in the first postnatal week, reaching definitive adult cell function and morphology quickly after the postnatal change in loading conditions and oxygenation^23,24^.

Our study aimed to be a proof of concept, that chronic perinatal hypoxia disrupts the process of normal cardiac development. To our knowledge, this is the first study to include both pre- and postnatal hypoxia to model the range of cardiac development affected by hypoxia in cyanotic CHD. Further, we incorporated a period of recovery to simulate definitive repair of cyanotic CHD. Limitations of our model include the inherent constraints of using small animals to model human disease, and the risk of introducing maternal stress into gestation. Litter size and pup size was reduced in our animals, which may be evidence for maternal stress. Indeed, maternal stress has emerged as an important predictor of poor outcomes in infants with CHD^50,51^.

Another limitation of our study was the inability to measure contractile function at P8 due to the small size of the hypoxic pups; thus precluding assessment of interval change following recovery. Phenotypic testing at P8 included sedated ECGs, and it is likely that hypoxic animals received a larger total dose of isoflurane to achieve adequate sedation. Importantly, the comparative effect of degree of hypoxia between species is unknown. We chose 10.5% for the degree of hypoxia, as a fractional inspired oxygen concentration (FiO_2_) of 10% correlates with pulse oximetry readings of 55-70% in rodents^52^. For comparison, human fetuses with cyanotic CHD have a mean oxygen saturation of 48% in the ascending aorta^7^, and neonates have target oxygen saturation ranges of 70-85%. Despite the unknowns between species, we believe this study is a first step toward understanding the impact of hypoxia on cardiac development.

The Cardiac Safety Research Consortium has implored the research community to perform more studies of developmental cardiac physiology to better understand the substrate on which cardiac therapies may work in the pediatric population^20^. Further studies regarding the effects of hypoxia on cardiac development may allow us to better target cardiac therapeutics for the cyanotic CHD population. A better understanding of the effects of chronic perinatal hypoxia on cardiac development could lead to improved surgical outcomes and overall improved cardiovascular health in the cyanotic CHD population.

## Acknowledgments

We gratefully acknowledge Dr. Susan Knoblach, Karuna Panchapakesan, and the Children’s National Research Institute Genomics and Bioinformatics Core for assistance with microarray experiments; we also acknowledge Dr. Norman Lee for assistance with gene expression and heatmap analysis, and Dr. Christopher Spurney and Dr. Charles Berul for helpful discussions.

## Author Contributions

JR, ZD, CM, DG, MR, LS, NV, MR performed experiments; JR, ZD, CM, DG, NGP analyzed data; JR, DG, NGP prepared figures; JR, DG, NGP drafted manuscript; JR, ZD, NI, NGP conceived and designed experiments; all authors approved manuscript.

